# Saltatory axonal conduction in the avian retina

**DOI:** 10.1101/2022.07.04.498722

**Authors:** Christoph T. Block, Malte T. Ahlers, Christian Puller, Max Manackin, Dipti R. Pradhan, Martin Greschner

## Abstract

In contrast to most parts of the vertebrate nervous system, the ganglion cell axons in the retina typically lack any myelination. Ganglion cell axons of most species only become myelinated once they leave the retina to form the optic nerve. The avian retina is a well known exception in that ganglion cell axons are partly myelinated in the retinal nerve fiber layer. However, the functional and structural properties of myelination in the nerve fiber layer remain elusive. Here, we used large-scale multi-electrode array recordings in combination with immunohistochemistry and fluorescence microscopy of European quail and pigeon retinas to investigate myelination of retinal ganglion cell axons. Intraretinal myelination was accompanied by the formation of nodes of Ranvier. The internode length was positively correlated with the axon diameter. The variability of internode lengths along each axon was significantly smaller than across axons. Saltatory conduction of action potentials was observed in a large population of recorded cells. On average, myelinated axons had higher conduction velocities than unmyelinated axons. However, both groups showed a significant overlap at low velocities. The number of simultaneously active nodes was positively correlated with the conduction velocity. In contrast, the internode length and the time it took a node to activate were weak predictors for the conduction velocity. However, the conduction velocity was well described by the number of activated nodes, the internode length, and the activation time in concert.

**Significance Statement:** Myelination of axons serves saltatory signal conduction, which greatly decreases the time it takes for an action potential to travel along an axon. Retinal ganglion cell (RGC) axons, as part of the central nervous system, are usually devoid of myelin in mammals, whereas avian RGC axons are myelinated well before they enter the optic nerve. Using high resolution multi electrode arrays, we were able to image the saltatory propagation of a spike across an axon. Most axons with saltatory conduction were faster than non-saltatory axons. Surprisingly, a large number of saltatory axons had low conduction velocities. The signal conduction patterns were more diverse than expected. The velocity could be explained by the number of simultaneously activated nodes of Ranvier, the internode length and their activation time.

## Introduction

Myelination of nerve cell axons is common in the vertebrate nervous system. Myelination is accompanied by clusters of voltage-gated ion channels in the membrane of axons, when gaps in the myelin sheath occur (reviewed in Van Hook et al., 2019). The periodic alternation of myelinated sections (i.e. internodes) with unmyelinated gaps where the cell membrane is exposed to the extracellular space (i.e. nodes of Ranvier), enables fast saltatory signal conduction. Since myelination increases the membrane’s length constant and decreases the time constant, it allows an activated node to depolarize subsequent nodes above threshold, avoiding time- and energy-consuming active spike generation in between, greatly increasing conduction velocity.

The healthy mammalian retina typically lacks any oligodendrocytes, which are the glial cells responsible for the formation of myelin sheaths wrapped around axons in other parts of the central nervous system (Gao et al., 2006). Since myelin is a lipid-rich, optically dense material it would potentially compromise the optical properties of the retinal nerve fiber layer, through which the light has to travel before reaching the photoreceptors. The axons of mammalian retinal ganglion cells (RGCs) therefore remain unmyelinated until they pass the lamina cribrosa to form the optic nerve, from where on myelination starts (Fujita et al., 2000). For mammals, a notable exception is the rabbit retina, where intraretinal myelination by oligodendrocytes is present, though limited to a certain region above the visual streak (Hughes, 1971; Schnitzer, 1985; Vaney, 1980). Intraretinal myelination has also been shown in fish, specifically in axons of presumed efferent origin (Witkovsky, 1971; Wolburg, 1980).

In contrast, oligodendrocytes are commonly found in the avian retina, where ganglion cell axons in the nerve fiber layer are myelinated accordingly (Imagawa et al., 1999; Inoue et al., 1980; Nakazawa et al., 1993; Seo et al., 2001; Won et al., 2000). The myelination is unevenly distributed across the retina, where axons are not as densely myelinated as they are in the optic nerve, and some axons remain unmyelinated altogether (Binggeli & Paule, 1969; Imagawa et al., 1999; Inoue et al., 1980). A distinguishing feature of myelin in avian retinas is its loose structure, characterized by largely missing intraperiod lines and proteolipid proteins, a major constituent of compact myelin in the central nervous system (Inoue et al., 1980; Kohsaka et al., 1980). In the optic nerve, only compact myelin is found (Fujita et al., 2001). Myelination of RGC axons in the nerve fiber layer is biased toward thicker axons (Nakazawa et al., 1993). Little is known, however, about how myelination is organized along individual RGC axons in the avian retina, and how it affects the conduction of spikes within the nerve fiber layer. If myelinated internodes alternate with clustered ion channels at nodes of Ranvier, as found in the optic nerve, one would expect a saltatory signal pattern, potentially in a subset of fast-conducting axons.

Here, we employed high-density multi-electrode arrays to record the electrical activity of RGC axons in quail and pigeon retinas, providing a functional analysis of saltatory spike conduction in the avian nerve fiber layer. Saltatory signal conduction was further explained by the localization of immunolabeled clusters of ion channels at nodes of Ranvier.

## Materials and Methods

### Animals and tissue preparation

All experiments were performed in accordance with the institutional guidelines for animal welfare and the laws on animal experimentation issued by the EU and the German government. Adult European quail (*Coturnix coturnix*) and pigeons (*Columba livia*) were killed by an overdose of pentobarbital (Narcoren, Boehringer Ingelheim). Adult C57BL/6J mice were killed by cervical dislocation. All animals were housed in a 12 hr light-dark cycle and experiments were performed during day hours.

### MEA recordings and analysis

Retinas were recorded as described previously (Field et al., 2007). Briefly, quail and pigeon retinas retinas were dissected under infrared illumination in Ringer’s solution (100 mM NaCl, 6 mM KCl, 1 mM CaCl2, 2 mM MgSO4, 1 mM NaH2PO4, 30 mM NaHCO3, 50 mM Glucose), bubbled with carbogen (95% O2 and 5% CO2), pH 7.5 (Stett et al., 2000) at room temperature. A 3 × 3 mm piece of pigment epithelium attached retina from the dorsal periphery was mounted, ganglion cell side down, on a large-scale CMOS (complementary metal-oxide-semiconductor) array (3Brain). MEA recordings were performed at 36.5°C in the chamber for the mouse and at 37.5°C for the avian retinas. Mouse retinas were dissected and recorded in carbogenated Ames’ medium (pH 7.4, Sigma).

Recordings were analyzed offline to isolate the spikes of different cells. After bandpass filtering (60 Hz - 4 kHz), candidate spike events were detected using a threshold on each electrode. The voltage waveforms on the electrode and neighboring electrodes around the time of the spike were extracted. Clusters of similar spike waveforms were identified as candidate neurons if they exhibited a refractory period. Duplicate recordings of the same cell were identified by temporal cross-correlation and matching electrical images and removed.

After spikes from different cells were segregated, the spike-triggered average electrical signal was calculated for each cell separately (Litke et al., 2004). Specifically, for each electrode, the voltage waveform during the time period from −1.62 ms to 4.42 ms around each spike was extracted. All spike waveforms were upsampled from 17.8 kHz to 200 kHz by cubic-spline-interpolation and aligned to the spike’s minimum. Waveforms were then averaged electrode-wise, temporally aligned to the occurrence of a somatic spike, to gain a spatio-temporal representation of a typical spike traveling along an axon. To construct the two dimensional electrical image, the minimum of the waveform at each electrode was displayed in the spatial layout of the array. If indicated in the figure legend, the soma amplitude was clipped to improve visibility. The electrical images were inspected for evidence of somatic and dendritic signatures and an axon of sufficient clarity and length. All back-propagating axons were excluded as the physiology of potential efferent axons would be questionable in an isolated piece of retina.

Axons were traced manually in the electrical images to adequately represent their structure. Positions of prominent local minima were chosen. To gain a signal profile along the axon the axonal trace was linearly interpolated from the manually defined positions. In the extracted signal profile, local minima and maxima were identified. The ratio of the robust means of all local minima and all local maxima, respectively, was calculated. A robust mean, i.e., the mean of the central 80% percentile, was chosen to exclude potential biases from border regions. The baseline of the entire dataset was adjusted to keep the index well behaved and in the range between 0 and 1. A low value of this continuity index represents a more discontinuous saltatory axonal signal profile, and a high value represents a more continuous non-saltatory signal profile. The conduction velocity of the spike was estimated by a linear regression of the euclidean distances between the identified positions on the axon and the time points of the corresponding minima of the signal waveform. The resulting slope is the conduction velocity of the spike. A Gaussian mixture model was applied to the continuity indices and the conduction velocities. On this basis, axons were separated into two groups of saltatory and non-saltatory spike conduction, respectively. RGCs with a likelihood < 0.95 to belong to either cluster were excluded from further analysis.

The prominent local minima in the electrical images of saltatory axons were defined as the position of nodes of Ranvier. The number of active nodes was defined as the number of orthodromic nodes with a simultaneously negative signal following a given active node. The robust mean across the nodes of each axon is reported. Node amplitude was defined as the minimum signal on the position of the nodes. The robust mean across the nodes of each axon is reported. Onset duration was defined as the time from half-minimum to minimum of the signal waveform at each node. The robust mean across the nodes of each axon is reported.

### Immunostaining and light microscopy

For immunostaining, the tissue was fixed in 4% paraformaldehyde in 0.01 M phosphate buffered saline (PBS, pH 7.4) for 20-30 minutes at room temperature. Immunostainings were performed on retinal tissue collected from enucleated eyes or on tissue which was mounted on multi-electrode arrays for recordings prior to the fixation. If the latter was the case, the tissue was carefully transferred from the array onto black cellulose filter paper (Millipore) prior to the fixation procedure. After fixation, the tissue was cryoprotected overnight with 30% sucrose in PBS and stored at −20°C until use.

Immunostaining was performed by an indirect fluorescence method. Retinal wholemounts were incubated at room temperature for 2-3 days in the primary antibody solution containing 5% normal donkey serum, 1% bovine serum albumin, 1% Triton X-100 and 0.02% sodium azide dissolved in PBS. Primary antibodies used in this study were anti-myelin basic protein (MBP, rat, monoclonal, clone 12, Bio-Rad, cat# MCA409S, 1:1000), anti-Neurofilament 200 (NF200, rabbit, polyclonal, Sigma-Aldrich, cat# N4142, 1:1000), and anti-voltage gated sodium channel, pan (panNav, mouse, monoclonal, clone K58/35, Sigma-Aldrich, cat# S8809, 1:1000). Secondary donkey antibodies (1:500, conjugated to Alexa 488, 568, 647, Invitrogen, or Cy3, Jackson ImmunoResearch) were incubated at room temperature for 4 hours in the same incubation solution. The tissue was mounted on glass slides and coverslipped with vectashield (Vector Laboratories). Spacers between glass slides and coverslips were used to avoid squeezing of the tissue.

Overview image stacks were taken with a confocal laser scanning microscope (Leica TCS SL), equipped with a 40x/NA1.25 oil immersion objective. For high resolution images of panNav clusters and corresponding size measurements, image stacks were acquired with a 63x/NA1.32 oil immersion objective. All confocal stacks were acquired with z-axis increments of 0.2 µm. Maximum intensity x/y- or z-projections are shown in all figure panels of microscopic images and were done in Fiji. Brightness and contrast of final images were adjusted in Adobe Photoshop. For measurements of internode lengths and axon diameters, tile scans of image stacks along axons were performed with a Leica DM6B fluorescence microscope, equipped with a motorized stage in combination with a 40x/NA1.3 oil immersion objective. Automated acquisition and stitching of individual tiles occurred through the Leica LASX software. Fiji was used to measure the distance between panNav clusters, i.e. nodes of Ranvier, along individual NF200-positive axons. Internodal lengths were measured with the segmented line tool as the distance between the centers of panNav-positive clusters, which allowed for precise tracing of axonal paths through the stitched image. Please note that our definition of internode lengths for the anatomical measurements includes the para- and juxtaparanodes, and is referring to the distance between panNav clusters. Each internode length measurement was accompanied by a measurement of axon diameter as an orthogonal line relative to the axon at the approximate half distance between a given pair of clusters. Measurements of panNav cluster lengths and diameters were performed in Fiji on high-resolution confocal image stacks acquired as mentioned above. A single image plane was manually selected per cluster to include the largest diameter of the whole tubular structure. A region of interest was cropped and automatically thresholded using Otsu’s method. The line tool was used in the thresholded image to measure the diameter and lengths. Individual lengths values represent the mean lengths averaged across measurements of both sidewalls of each cluster.

### Statistical analyses

For the correlation coefficient the lower and upper bounds of a 95% confidence interval were reported. Statistical tests for zero correlation with conduction velocity were rejected for all investigated parameters. The test for internode length versus conduction velocity led to p<0.05. The test for onset duration versus conduction velocity led to p<0.01 and all other to p<0.001. For the analysis of internode length variation within and across axons (Figure 7E) the quartile coefficient of dispersion was used (Bonnett, 2006). The coefficient of dispersion across axons was bootstrapped with replacements. A Wilcoxon rank sum test rejected equal medians of these distributions (p<0.001). In the results section, all measurements are reported as the mean ± SD. Outliers were included in the statistical analysis. To address the variability in raw measurements, a robust mean on the central 80% of data was used (see above). The analysis was performed in Matlab (MathWorks).

## Results

### Electrical images reveal saltatory spike conduction

The high electrode density of large scale multielectrode arrays enables the analysis of spatio-temporal electrical activity of the recorded neurons (Greschner et al., 2014; Litke et al., 2004; Radivojevic et al., 2017; Zeck et al., 2011). The electrode-wise spike-triggered average electrical activity over time reveals how a typical spike travels along the axon of a retinal ganglion cell (RGC). Collapsing the time dimension by a minimum projection makes the spatial structure of the electrical activity of the recorded cell apparent, i.e., the electrical image. The electrical images of the European quail RGCs showed the expected spatial structure, i.e., a high-amplitude somatic area, a surrounding dendritic area, and an axon extending toward the optic disk (Figure 1). For many of the recorded RGCs we found a continuous axonal conduction (Figure 1 A-C), as is common for unmyelinated RGC axons in most mammalian species. However, a larger fraction of the recorded quail RGCs showed a discontinuous, saltatory axonal spike conduction (Figure 1 D-F, see supplementary movie S1). In those cases, we found regions of high amplitude signals that were interchanged with lower-amplitude regions of opposite signal polarity, as expected for myelinated axons with passively conducting regions, periodically interrupted by active node-of-Ranvier-like structures (Fig 1D, F). Analysis of the spike conduction along the axon over time revealed a roughly linear distance-time relation in both cases (Fig 1B, E). The slope of a linear fit of the times of maximum negative deflection and the corresponding distances along the axon was used as an estimate of the axonal conduction velocity.

**Figure 1:**
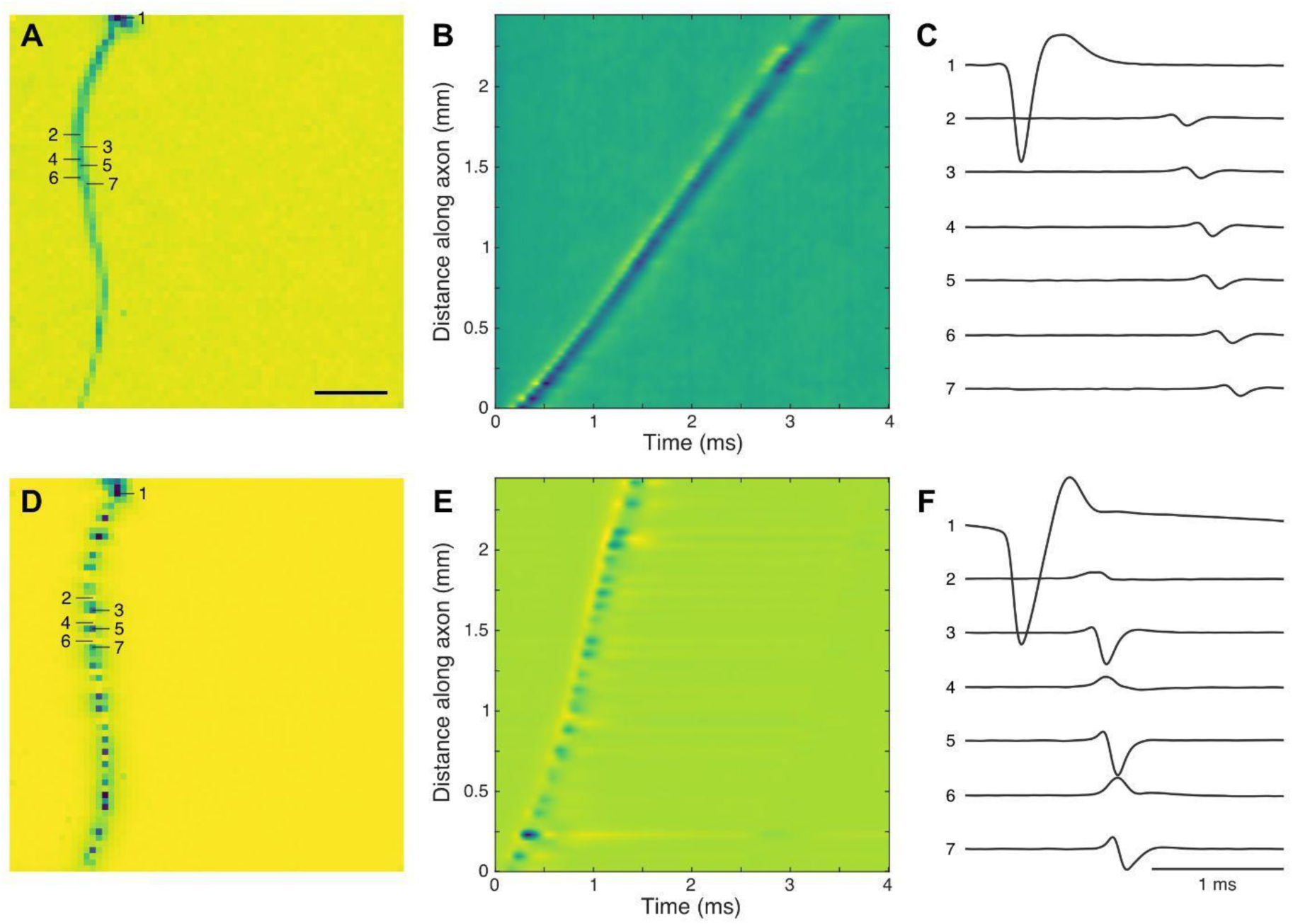
The electrical images of European quail retinal ganglion cells revealed axons with and without saltatory conduction. **A:** Electrical image of non-saltatory RGC with conduction velocity v = 0.8 m/s. Minimum voltage projection along the time dimension of the spike triggered average electrical activity. Colder colors indicate more negative potentials. 64 × 64 electrodes, Scale bar: 500 µm **B:** Signal waveform along the axon. Colder colors indicate more negative potentials. **C:** Signal waveform for seven selected electrodes indicated in A. Scale bar: 1 ms. Y-axis in arbitrary units, same y-scale in C and F. **D-F:** As A-C for saltatory RGC with v = 2.0 m/s.

We quantified the degree of signal discontinuity of each traced axon by calculating the ratio between the local maxima and minima along the axon in the electrical image. The distribution of this ratio was bimodal (Figure 2), reflecting the differentiation of axons into saltatory and non-saltatory conduction. The 42 µm electrode-to-electrode distance of the multielectrode array did not allow for reliable detection of the smallest internode lengths. This biased our functional classification toward RGCs with larger internode lengths and potentially misclassified axons with small internode lengths as non-slatatoric axons (see methods for more details). Thus, for further analysis, we excluded neurons with a low likelihood to belong to either cluster (Figure 2, gray cluster). As expected, saltatory axons displayed an overall higher conduction velocity (saltatory: 1.24 ± 0.35 m/s, non-saltatory: 0.71 ± 0.12 m/s). While lower conduction velocities were found in both non-saltatory and saltatory axons, high conduction velocities were only found in saltatory axons.

**Figure 2:**
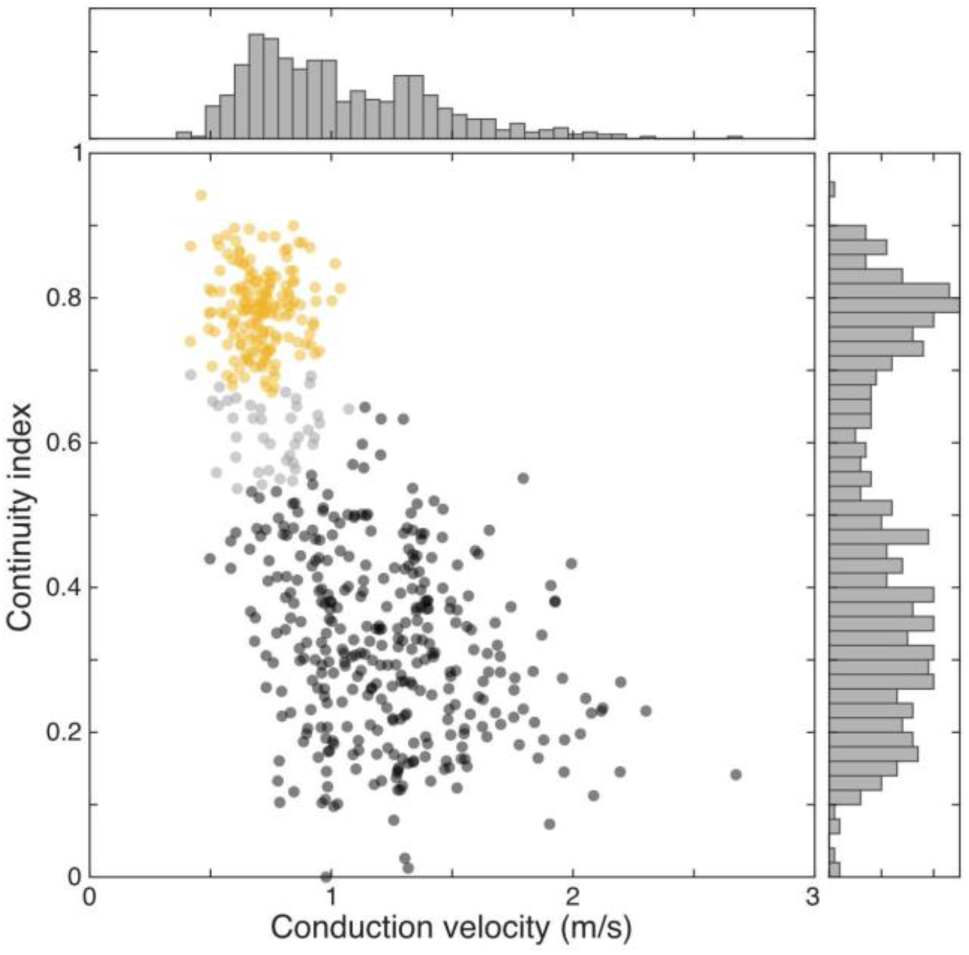
Saltatory and non-saltatory axons showed different conduction velocities. The RGCs with a sufficient axon length of one European quail retina were classified into 333 saltatory (black) and 164 non-saltatory (yellow) axons. The overlap region of 40 axons (gray) was excluded from further analyses (likelihood < 0.95, mixture of Gaussians). The continuity index is the mean peak amplitudes divided by the mean trough of amplitudes along the axon. See methods for details. Figures 3-6 present data from the same cells. See supplementary material for a second quail retina (S2).

**Figure 3:**
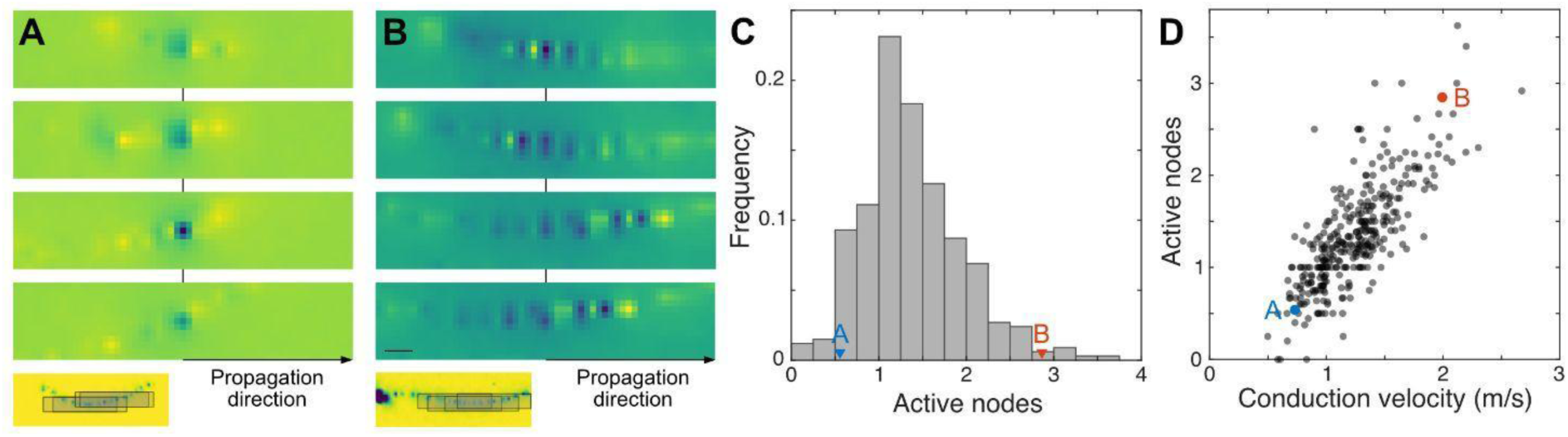
Number of active nodes was correlated with conduction velocity. **A:** Enlarged view of spatial electrical activity of four example nodes from one RGC axon. Center in space and time was defined as the local minimum of the electrical signals. Inset illustrates the spatial origins of the enlarged views. Blue colors indicate negative values. Scale bar = 100µm. **B:** As A for a second RGC with a high number of active nodes. **C:** Histogram of number of active nodes, defined as the robust mean of the number of counted nodes in propagation direction. RGCs of A and B marked. **D:** Number of active nodes against conduction velocity. RGCs of A and B marked.

**Figure 4:**
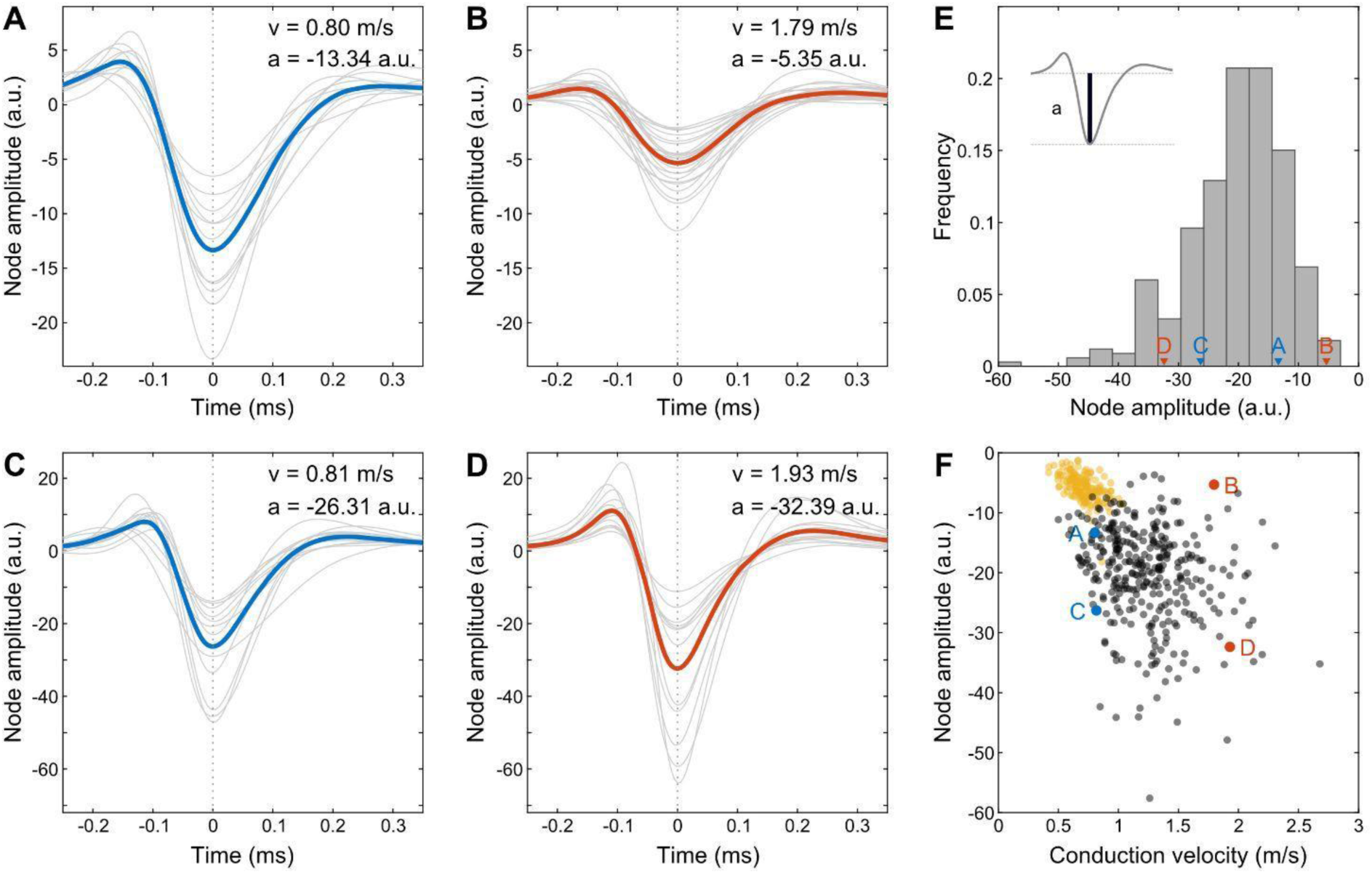
The node amplitude showed a large variation across axons. **A-D:** Node amplitudes of saltatory axon segments of four different RGCs. Gray lines represent individual nodes, and the bold line represents the mean. Arrangement as in F. **E:** Histogram of number of node amplitudes across axons. RGCs of A-D marked. Slower (blue) and faster (red) axons marked for visual guidance. **F:** Node amplitude against conduction velocity for saltatory (black) and non-saltatory axons (yellow).

### Large variability of biophysical parameters

Considering the factors that influence the axonal conduction velocity, the number of orthodromically active nodes, i.e., nodes simultaneously active in the spike propagation direction, could be expected to play a dominant role. These indicate how far the signal on the node with currently the largest signal amplitude is spreading. The number of active nodes in propagation direction was on average 1.35 ± 0.75 and indeed correlated with the conduction velocity (correlation coefficient: 0.78; CI: [0.74, 0.82]). In addition, the number of still active antidromic nodes was correlated with the number of active orthodromic nodes (correlation coefficient: 0.61, CI: [0.54,0.67]). Leading to a total spatial extent of a spike of 3.06 ± 1.00 (minimum: 1, maximum: 7.13) nodes.

A parameter determining the number of orthodromically active nodes is the length constant of the axon which is mainly influenced by the diameter of the axon and its level of myelination. In addition, the number of active nodes is influenced by their overall sensitivity, i.e., how readily nodes can be activated by the signal spread from a preceding node. One way to approach this in our data was the measurement of the signal amplitude at the nodes. While the conduction velocity roughly scaled with the signal amplitude for the non-saltatory axons (correlation coefficient: −0.53, CI: [−0.64, −0.42]), this was less apparent for the saltatory axons (correlation coefficient: −0.23, CI: [−0.33, −0.13]).

Besides the number of orthodromic nodes the signal spreads to, the internode length is expected to be a major factor for the conduction velocity (Brill et al., 1977). However, no correlation between the internode length and conduction velocity was apparent (Figure 5, correlation coefficient: 0.14, CI: [0.03, 0.24]). Axons with the same internode length had widely different conduction velocities and axons with the same conduction velocity had widely different internode lengths.

**Figure 5:**
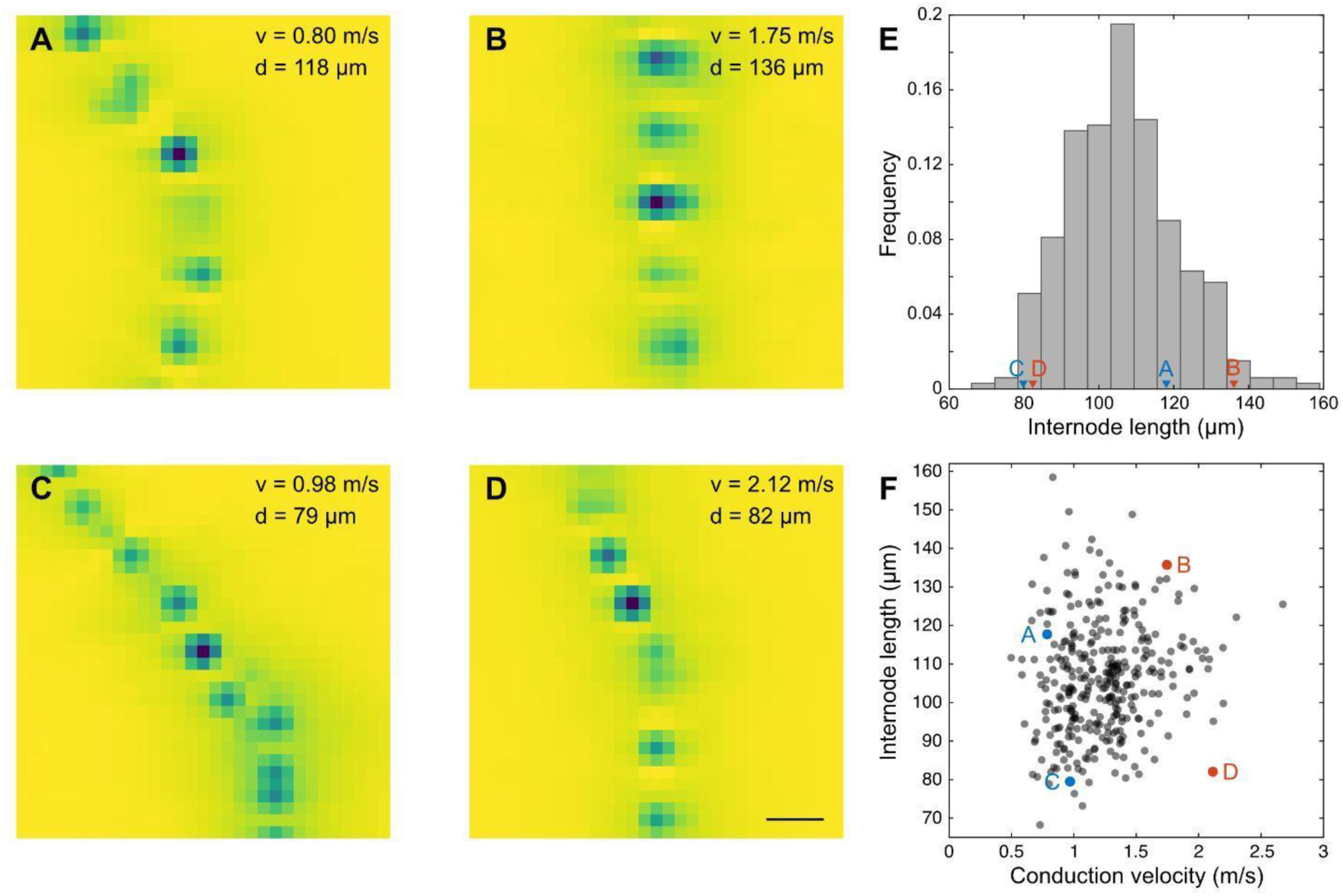
The internode length showed a large variation across axons. **A-D:** Electrical images of four saltatory ganglion cell axon segments. Arrangement as in F. Blue colors indicate negative values. Scale bar = 100µm. **E:** Histogram of number of internode lengths. RGCs of A-D marked. **F:** Internode length against conduction velocity. Please note that the electrode-to-electrode distance of the multielectrode array of 42 µm is too large to reliably measure the small end of the internode lengths (see methods).

In addition to the distance a signal is able to spread, the time it takes to activate a given node is a major determinant of the conduction velocity. However, the signal onset of a given node showed no apparent correlation with the conduction velocity (Figure 6, correlation coefficient: −0.15, CI: [−0.25, −0.04]).

**Figure 6:**
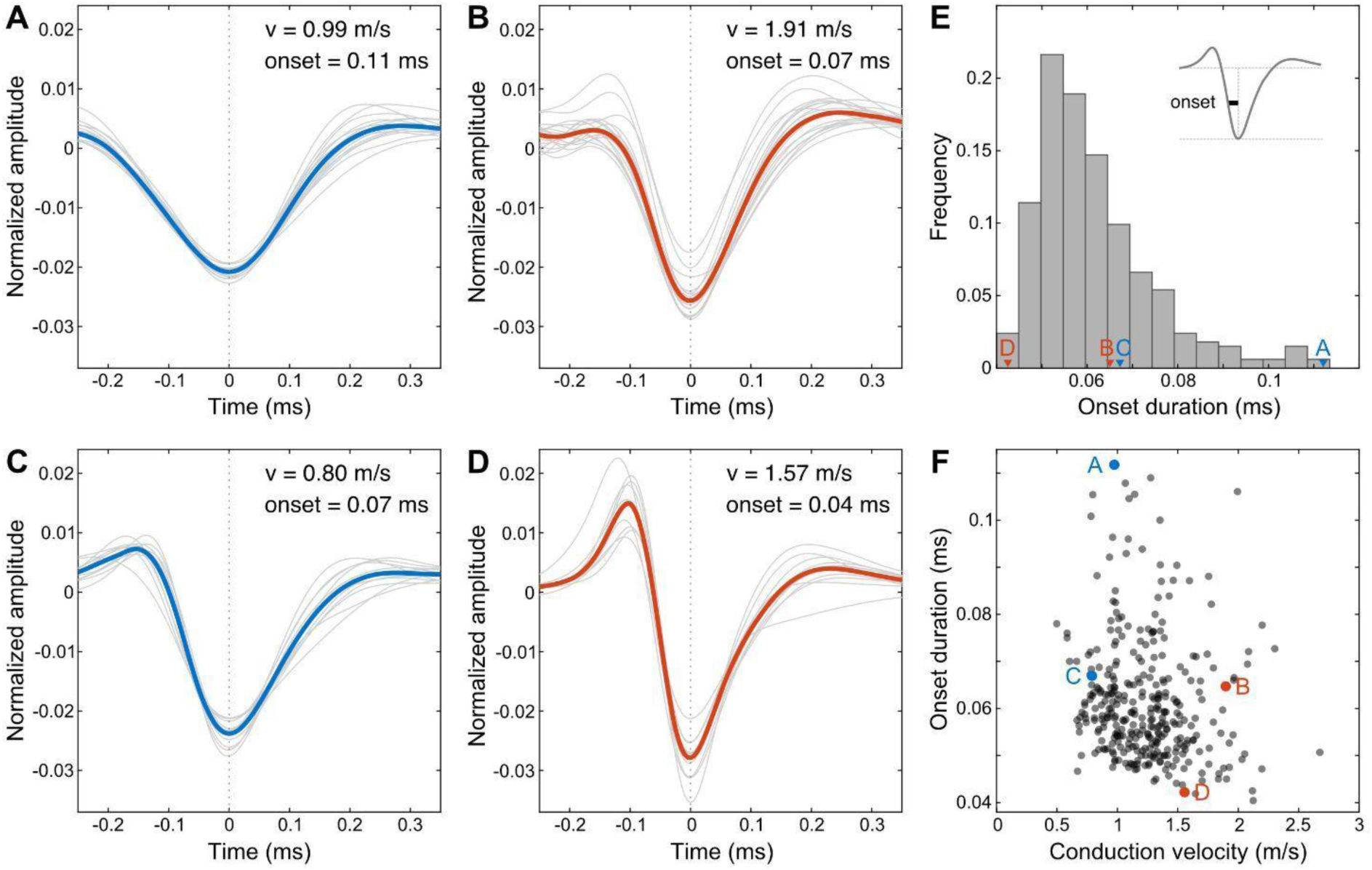
The onset duration showed a large variation across axons. **A-D:** Spatial electrical activity of four saltatory ganglion cell axon segments. Gray lines represent individual nodes (normalized to absolute area) and the bold lines represent the mean. Arrangement as in F. **E:** Histogram of onset durations across axons. RGCs of A-D marked. **F:** Internode length against onset duration.

While the individual node properties investigated above (amplitude, internode length, onset duration) did not have an obvious effect on signal conduction velocity a descriptive model of the number of active nodes multiplied by the internode length, divided by the onset time of the activation on a given node predicted the conduction velocity reasonably well (correlation coefficient: 0.83, CI: [0.79, 0.86]).

### Voltage-gated sodium channels are clustered at nodes

Figure 7A-C shows the pattern of myelination in the temporal region of a quail retina where MEA recordings were typically performed. Sodium channel clusters occurred as short, tubular structures and were only found at myelinated axons. Similar to conventional nodes of Ranvier, no myelination was found at or in close vicinity to the clusters. These clusters were never observed at axons where myelin was absent. Unmyelinated axons showed only diffuse panNav immunoreactivity which rarely exceeded background levels. Thus, those axons are not represented in figure 7C, as they were often negative for the neurofilament subunit labeled here (see Methods). The expression levels of neurofilament differed between individual axons. Myelination and the corresponding clustering of sodium channels occurred at both strongly and weakly neurofilament-positive axons. The sodium channel cluster length was 1.4 ± 0.3 μm and the cluster width was 1.1 ± 0.3 μm (n = 52). The distances between sodium channel clusters were measured along strongly neurofilament-positive axons (Figure 7D). This internode length was significantly more variable across axons than along individual axons (Figure 7D, Wilcoxon rank-sum test: p<0.001). We used the width of the neurofilament labeling in the middle between each pair of clusters as an approximation of the axon diameter (Figure 7F). The internode length (mean: 76.8 ± 25.2 µm, n = 200) was positively correlated with the axon diameter (mean: 0.9 ± 0.2 µm, n = 200, correlation coefficient: 0.55, CI: [0.44, 0.64]).

### High conduction velocities in the pigeon retina

Results of experiments with pigeon retina were largely consistent with those obtained from the quail. panNav-immunoreactive nodes were located along neurofilament-positive axons (Figure 8A). Accordingly, electrical images revealed saltatory signal conduction, alongside those with no obvious saltatory features (Figure 8B, C). Remarkably, spike conduction observed in the pigeon retina was not only substantially faster (2.0 ± 0.73 m/s) than in those of the European quail (0.91 ± 0.34 m/s), also the spread of conduction velocities was considerably broader in pigeon axons. Conduction velocities in mouse retinas were on average slower (0.65 ± 0.15 m/s) than those of the considered avian species (Figure 8D).

**Figure 7:**
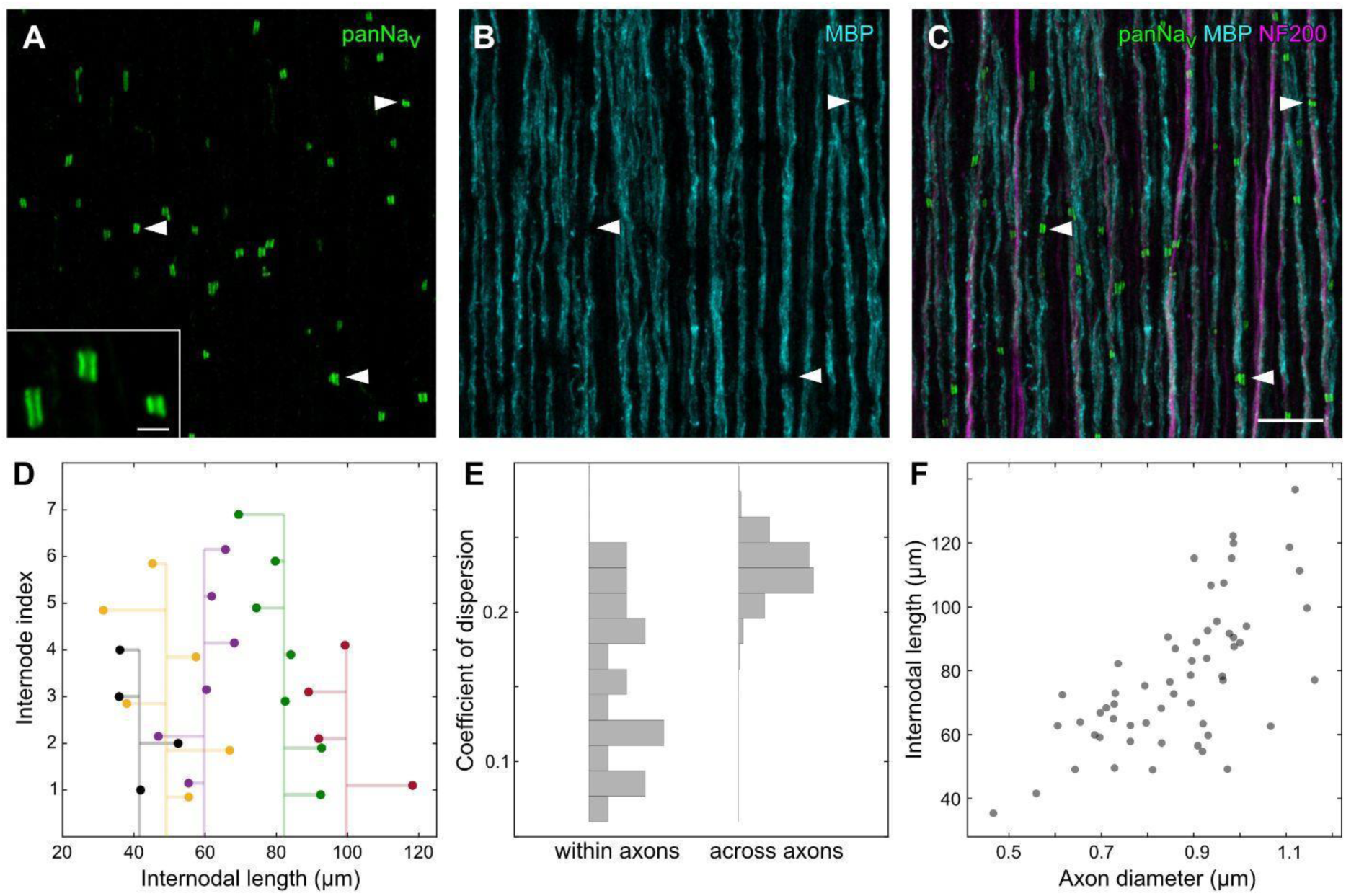
Voltage-gated sodium channels were clustered at nodes of myelinated ganglion cell axons in the quail retina. **A-C:** Confocal images of the nerve fiber layer in a quail retina immunolabeled against voltage-gated sodium channels (panNav), myelin basic protein (MBP), and neurofilament (NF200). Dense clusters of Nav channels were found along axons at sites where myelination was interrupted (example positions marked by arrowheads). Inset in A shows panNav clusters at higher magnification. Scale bars, C: 10 µm, inset in A: 2 µm **D:** Successive internode lengths along five example axons. Vertical line represents the mean internode length for a given axon. **E:** The variation of internode lengths is significantly smaller within an axon than across axons (quartile coefficient of dispersion). Subselection of axons with at least five measured nodes (n = 22). Variance across axons is bootstrapped. **F:** The distances between nodes at axons increase as a function of axon diameters. Mean over repeated measurements along individual axons (n = 57).

**Figure 8:**
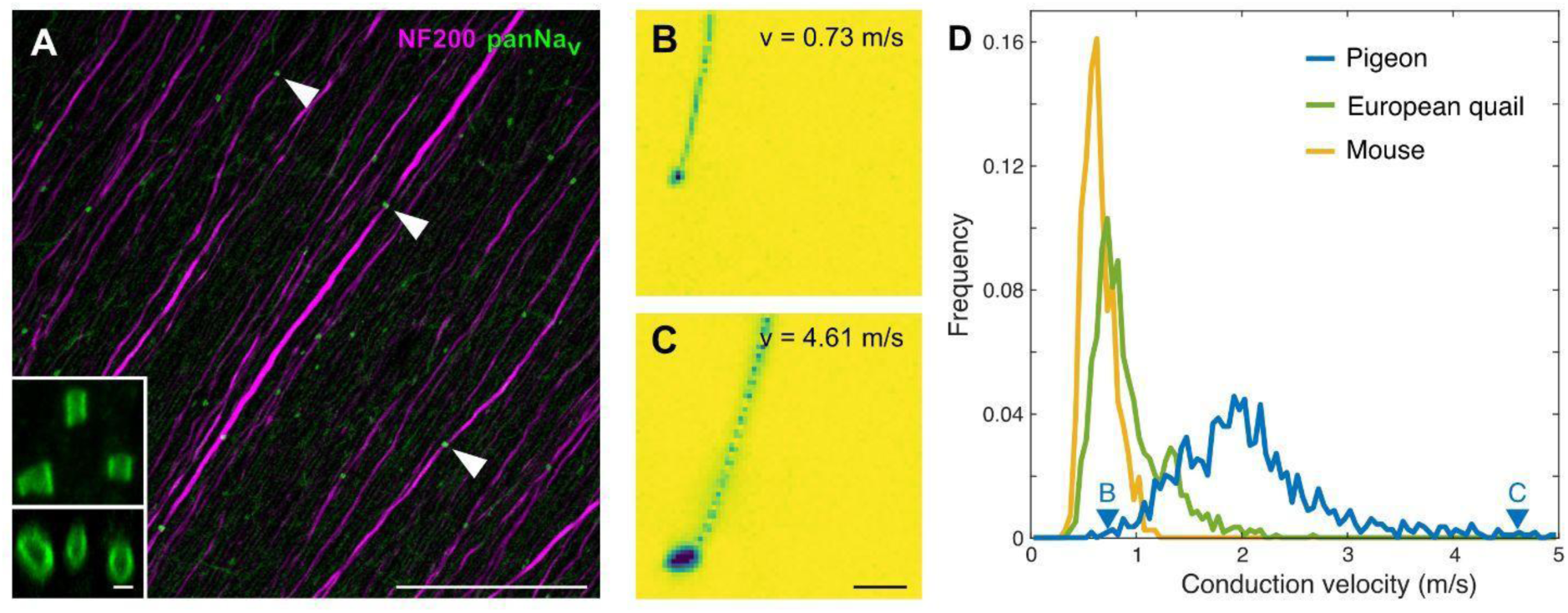
Saltatory conduction and myelination in the pigeon retina. **A:** Confocal images of the nerve fiber layer in a pigeon retina immunolabeled against voltage-gated sodium channels (panNav), and neurofilament (NF200). As in quail retina, clusters of Nav channels are found on axons (example positions marked by arrowheads). Inset in A shows panNav clusters at higher magnification: Top: maximum intensity projection of an image stack. Bottom: side view of the rotated image stack. Scale bars: 50 µm, inset: 1 µm **B, C:** Example electrical images of slow (B) and fast (C) spike conduction in pigeon RGCs, scale bar: 500 µm. See supplementary movie S1. **D:** Conduction velocity histogram for pigeon (blue, 2 retinas, 1141 axons), quail (green, 2 retinas, 1175 axons), and mouse (yellow, 2 retinas, 410 axons). Histograms were normalized to the total number of axons analyzed for the respective species. Blue arrowheads indicate velocities of cells shown in B and C. Please note that the recordings were performed below body temperature of the animals.

## Discussion

We imaged saltatory conduction in the European quail and pigeon retina and confirmed intraretinal myelination and the formation of nodes of Ranvier. Two subpopulations of the recorded cells were classified as saltatory or non-saltatory, respectively, according to their axonal signal profile in the electrical image. On average, saltatory axons exhibited higher conduction velocities. However, a strong overlap of the conduction velocities between both groups was apparent for lower velocities. The number of simultaneously active nodes was positively correlated with the conduction velocity. In contrast, the internode length and the time it took a node to activate were only weak predictors for the conduction velocity. However, the conduction velocity was well predicted by the number of activated nodes, the internode length, and the activation time in concert.

All morphological features of the panNav-positive sodium channel clusters at axons in quail and pigeon retinas, i.e. their tubular shape and size, their distribution pattern along axons and the myelin-sheath gaps at those sites, closely resembled the structures shown for conventional nodes of Ranvier in the vertebrate optic nerve or elsewhere in the central nervous system (Arancibia-Cárcamo et al., 2017; Boiko et al., 2001; Caldwell et al., 2000; O’Brien et al., 2008; Rasband et al., 1999). Here, we have used a pan antibody directed against a highly conserved sequence present in all vertebrate Nav1 isoforms (Rasband et al., 1999). Therefore, the exact sodium channel subunit expressed at nodes of Ranvier in quail and pigeon RGC axons remains unknown. In the mammalian retina, RGCs express Nav1.6 at the axon initial segment and Nav1.2 in their unmyelinated axons within the nerve fiber layer. There, nodes of Ranvier are only formed in the optic nerve and Nav1.6 typically replaces Nav1.2 during development. Nav1.8 may represent an additional candidate for nodes of alpha-like cell types with large-caliber axons (reviewed in Van Hook et al., 2019). Thus, it is tempting to speculate that, in birds, the diffuse panNav labeling observed in unmyelinated axons represents Nav1.2, while the clustered channels at nodes of Ranvier are formed mostly by Nav1.6, with Nav1.8 expressed only by a subset of particularly thick axons. However, further studies are required to reveal the identity of voltage-gated sodium channel subunits and their individual kinetics in RGC axons of the avian retina.

Overall, our functional results for the retina are consistent with data for other parts of the nervous system. However, specific constraints influence the axonal conduction in the retina. The need for good optical transparency of the nerve fiber layer limits the structure and amount of myelin. Myelin acts as an electrical insulator that, by reducing the internodal membrane conductance and capacitance, increases the length-constant, decreases the time-constant, and ultimately raises the conduction velocity of the axon. Thus, a balance between improvement in conduction velocity and impairment of optical quality of the nerve fiber layer is likely for intraretinal axons.

In fact, only a subgroup of the intraretinal RGC axons is myelinated (Imagawa et al., 1999; Inoue et al., 1980). It is easy to speculate that RGC types that transmit information related to alertness or motion processing are privileged by myelination and fast conduction velocities. In addition, all efferent axons are described to be myelinated intraretinally (Wilson & Lindstrom, 2011). We observed a surprising overlap in conduction velocities between saltatory and non-saltatory axons, specifically, myelinated, low-velocity axons were found. Myelination increases the conduction velocity of an axon linearly with its diameter, while the velocity in unmyelinated axons increases only with the square root of the diameter (Brill et al., 1977; Herman, 2007; Waxman, 1980). Myelinated axons are expected to reach higher velocities than unmyelinated axons for diameters larger than 0.2 µm (Waxman & Bennett, 1972). Thus, the observed low velocities of saltatory, i.e. myelinated axons could have as well been achieved by unmyelinated axons of comparable diameter, gaining a supposedly better transparency. The potential of higher spatial packaging density of myelinated axons also offers no plausible explanation, as the total diameter of an axon including myelin can be expected to be larger than that of an unmyelinated axon for low velocities (Waxman, 1980). Also, we found no obvious difference in the firing rates of myelinated and unmyelinated axons of similar velocity, thus giving no hint toward a potential metabolic optimization by myelination.

The intraretinal conduction velocities were relatively low compared to other parts of the central nervous system. For example, cortico-spinal neurons can reach conduction velocities of 100 m/s (cat, Takahashi, 1965), thalamic-cortical conduction velocities are in the range of 6-38 m/s (rabbit, Swadlow & Weyand, 1985) but also slow cortical neurons with velocities as low as 0.3 m/s were described (rabbit, Swadlow, 1990). Extraretinal conduction velocities of RGC axons are indeed increased in comparison to the intraretinal velocities (Stanford, 1987; Stone & Freeman, 1971), making it likely that a fast conduction is generally beneficial and further substantiating the notion of constrained intraretinal myelination.

The conduction velocity had no apparent correlation with the internode length in our data. Theoretical studies predicted a maximized conduction velocity for a relatively wide range of internode lengths of 100-200 times the axon diameter (Brill et al., 1977; Moore et al., 1978). The reported ratio of internode length to axonal diameter fits our observation for the retina. Additionally, while at first glance, the variation of measured internode lengths along individual axons might be surprising, similar variations have been described in regions of the peripheral and central nervous systems of different vertebrates (chicken auditory brainstem: Seidl et al., 2010; zebrafish spinal cord: Auer et al., 2018, Vagionitis et al., 2022; rabbit spinal cord: Hess & Young, 1949; rabbit peripheral nerves: Vizoso & Young, 1948).

Considering the internode lengths reported here, it has to be noted that the detection of nodes in the physiological data was limited by the electrode-to-electrode distance of the multielectrode array. As expected by the sampling theorem, internode lengths below ∼90 µm could not be reliably detected and our analyses thus underrepresented RGCs with smaller internode lengths. On the other hand, the anatomical analysis was biased toward smaller internode lengths as RGC axons could only be reliably traced over shorter distances based on the neurofilament staining. Another methodological bias concerns the temperature of the retinas during recordings. While the temperature was relatively high for common standards of physiological recordings, it was still below the body temperatures of the respective animals. In particular, for the comparison of the conduction velocities of mouse, quail, and pigeon the relative temperature bias was larger for the avian species. The conduction velocity is expected to increase by roughly 50% over 10°C (reviewed in Waxman, 1980). Therefore, the conduction velocities should not be considered in absolute terms.

We observed significantly larger conduction velocities in the pigeon retina than in quail and mouse. It is critical for the fast flying pigeon to process visual information quickly (Boström et al., 2016; Potier et al., 2020). For example, the observed velocity difference of around 4 m/s between slow and fast axons in the pigeon retina (Figure 8 B-D) would lead to a difference in conduction delays of 12 milliseconds and a corresponding flight distance of approximately 0.3 m. The avian retina is a good example for the trade-off between transparency of the nerve fiber layer and fast intraretinal signal conduction, resulting in a large variety of biophysical parameters across RGCs axons.

## Acknowledgements

Research was supported by SFB 1372, Deutsche Forschungsgemeinschaft (MG) and RTG 1885/2, Deutsche Forschungsgemeinschaft (MG).

